# BMP2 binds non-specifically to PEG-passivated biomaterials and induces substantial signaling

**DOI:** 10.1101/2024.03.14.585026

**Authors:** Jean Le Pennec, Amaury Guibert, Romain R. Vivès, Elisa Migliorini

**Author notes:** Co-Corresponding authors Migliorini Elisa (E.M.), Romain Vivès (R.R.V.).

## Abstract

Biomaterials are widely employed across diverse biomedical applications and represent an attractive strategy to explore physiologically how extracellular matrix components influence the cellular response. In this study, we aimed to use previously developed biomimetic streptavidin platforms to investigate the role of glycosaminoglycans (GAGs) in bone morphogenetic protein 2 (BMP2) signaling. However, we observed that the interpretation of our findings was skewed due to the GAG-unrelated, non-specific adsorption of BMP2 on components of our biomaterials. Non-specific adsorption of proteins is a recurrent and challenging issue for biomaterial studies. Despite the initial incorporation of anti-fouling poly(ethylene glycol) (PEG) chains within our biomaterials, the residual non-specific BMP2 adsorption still triggered BMP2 signaling within the same range as our conditions of interest. To tackle this issue, we explored various options to prevent BMP2 non-specific adsorption. Specifically, we tested alternative constructions of our biomaterials on gold or glass substrate using distinct PEG-based linkers. We identified the aggregation of BMP2 at neutral pH as a potential cause of non-specific adsorption and thus determined specific buffer conditions to prevent it. We also investigated the induced BMP2 signaling over different culture periods. Nevertheless, none of these options resulted in a viable suitable solution to reduce the non-specific BMP2 signaling.

Next, we studied the effect of various blocking strategies. We identified a blocking condition involving a combination of bovine serum albumin and trehalose that successfully reduced the unspecific attachment of BMP2 and the non-specific signaling. Furthermore, the effect of this blocking step was improved when using gold platforms instead of glass, particularly with Chinese hamster ovary (CHO) cells that seemed less responsive to non-specifically bound BMP2 than C2C12 cells.

## Introduction

Biomaterials are commonly used for many applications, encompassing biosensors, drug delivery systems, biomedical implants, and the exploration of cellular responses in a more physiological environment than classically used plastic substrates. In this context, the non-specific adsorption of molecules or bacteria constitutes a significant issue hampering the development of new applications^1, 2^. The underlying mechanisms of protein non-specific binding predominantly entail interactions between hydrophobic domains and surfaces, potentially augmented by electrostatic interactions^3^. To tackle this issue, common anti-fouling strategies include the functionalization at surfaces of highly hydrophilic components with a neutral charge, providing a barrier to protein adsorption^4-6^. In particular, poly-ethylene glycols (PEG) synthetic polymers are commonly employed^5-7^. The reduction of protein adsorption exerted by PEGs can be explained physically by the generated steric hindrance between the protein and the surface, and chemically by the strong interaction between the PEG ether linkage and water molecules, which can hardly be overcome by proteins, thereby reducing their adsorption on surfaces. The PEG grafting density and polymer length also modify the conformation of the coating, but it has an unclear role in its anti-fooling performance^8-10^. Despite having a well-established role in reducing non-specific protein adsorption, common coating strategies, including PEGs, are not entirely satisfying, and their performance is protein-dependent^9^.

In previous studies, we developed PEG-based biomaterials (streptavidin biomimetic platforms) to facilitate the investigation of the roles of extracellular matrix components in cellular responses under physiological conditions^11-14^. In particular, we have been interested in investigating the roles of different glycosaminoglycans (GAGs) in the regulation of bone morphogenetic protein 2 (BMP2) signaling^15^. Developments were realized to functionalize the biomimetic platforms on glass or gold substrates with PLL-g-PEG-biotin^16^ or thiol-PEG-biotin linkers^11, 13^, ensuring certain surface passivation. These biomaterials were also designed to be compatible with microscopy readouts developed for high-content applications^16, 17^. We have commonly assessed the non-specific adsorption of BMP2 (on other components than GAGs) *ex-situ* with a quartz crystal microbalance (QCM) monitoring technique, revealing relatively low levels in comparison to specific binding on GAGs^16^.

Existing studies on BMP2 adsorption have predominantly focused on hydrophobic substrates utilized in biomedical implant applications. These investigations have primarily examined the ability of substrates to bind and release BMP2 while preventing protein unfolding that could hinder its bioactivity^18-24^. To our knowledge, concerns regarding non-specific BMP2 adsorption have not emerged as long as bioactivity is maintained for biomedical applications.

The present study highlights that even negligible amounts of non-specifically adsorbed proteins can induce a significant cellular response, complicating the interpretation of experimental outcomes of *in vitro* conditions. Specifically, the non-specific adsorption of BMP2 hindered our ability to elucidate the distinctive roles of various GAGs in the modulation of BMP2 signaling. We examined the effects of BMP2 adsorption on gold or glass biomaterials at different time points and with other proteins of the BMP family (BMP4 and BMP7). To circumvent this adsorption issue, we investigated the aggregation of BMP2 in different buffers that could be involved in these adsorption mechanisms. The quantification of BMP2 adsorption onto distinct components of our biomaterials enabled us to determine whether this adsorption was specific to these elements. Finally, we explored various blocking strategies to reduce BMP2 adsorption and the associated non-specific signaling in two cell lines on gold or glass platforms to identify specific conditions compatible with our *in vitro* cellular studies.

## Material and methods

### Buffers and Molecules

Unless stated otherwise, a solution termed Hepes, containing 10 mM Hepes and 150 mM NaCl (Sigma-Aldrich) at pH 7.4, was utilized for dilution and rinsing samples. Phosphate buffer saline (PBS, Sigma-Aldrich) was adjusted at pH 7.4 if necessary, and a sodium acetate (Sigma-Aldrich) buffer was prepared at 10 mM with a pH of 4.2. All buffers supplemented with 0.02% Tween-20 are consistently termed with the –T suffix (Hepes-T, PBS-T, Acetate-T). Biomimetic platforms were prepared by functionalizing mPEG-Thiol and biotin-PEG-Thiol (Polypure) on a gold substrate, or PLL(20)-g[3.5]-PEG(2)/PEGbiotin(3.4)50% linker (PLLgPEG, SuSoS AG) for glass substrates. Platforms were further conjugated with streptavidin (SAv, 55 kDa, Sigma-Aldrich). Details about biotinylated cyclic arginyl-glycyl-aspartic acid (RGD, 3.9 kDa) can be found in prior works^13, 14^. Heparan sulfate (HS) sourced from porcine intestinal mucosa was acquired from Celsus Laboratories. It has an average molecular weight of 12 kDa and a polydispersity of 1.59^25^. Chondroitin Sulfate A (C9819, CS-A, bovine trachea, average molecular mass 28 kDa^26^) and Dermatan Sulfate (C3788, DS/CS-B, porcine intestinal mucosa) were purchased from Sigma-Aldrich. Chondroitin Sulfate D (400676, CS-D, shark cartilage, average molecular mass of 38 kDa^27^) was acquired from Seikagaku. Chondroitin Sulfate E (CS-E) and Hyaluronic Acid (HA) were gifts from Kawthar Bouchemal. BMP2 produced was acquired from Medtronic (26 kDa, InductOs). Both BMP4 and BMP7 were purchased from R&D Systems.

### Dynamic light scattering BMP2 size measurement

The size distribution by volume of BMP2 aggregates in different media was determined by dynamic light scattering (DLS; Zetasizer Ultra, Malvern Panalytical, Worcestershire, United Kingdom). Light scattering was measured in a low-volume disposable sizing cell (ZSU1002, Malvern) at 25°C after an equilibration time of 300 s. The DLS size distribution was calculated from the autocorrelation curves applying the protein analysis algorithm (“general purpose” option) of the Zetasizer Software 7.11 (Malvern). A refractive index (RI) of 1.45 and an absorption coefficient of 0.001 were defined for BMP2, using the RI of water as solvent (RI: 1.33). Before measurement, BMP2 was stocked at 1 mg/mL in HCl 1mM (pH 3.0). The stock aliquot was centrifuged for 10 min at 13200 rpm preceding its dilution at 100 µg/mL with different buffers in low-binding tubes. Samples were briefly centrifuged for 90 s before being placed in measurement cuvettes. Triplicate measurements were performed for each sample. In the case of the “weak scattering” indication provided by the software data quality guidance, measurements were removed from the analysis.

### Biomimetic platform’s surface functionalization

Glass coverslips (24×24 mm; Menzel Gläser) underwent a gold-sputtering process (1.5 nm Cr and 8 nm Au) using a Plassys™ evaporating machine in a clean room, constituting the base substrate of gold biomimetic platforms. The gold-sputtered surfaces were activated with UV/Ozone Procleaner™ (BioForce Nanosciences) for 10 min and immersed overnight in an ethanol solution with 0.95 mM mPEG-thiol and 0.05 mM biotin-PEG-thiol before blow-drying the surfaces with nitrogen. SAv, cRGD, HS, and BMP2 were incubated sequentially by a liquid-handling robot (Evo 100, Tecan), performing the intermediate rinsing steps^16^. SAv was incubated at 10 µg/mL for 30 min to construct a monolayer. Then, co-functionalization of cRGD and GAGs was performed with a 10-minute incubation of biotinylated cRGD at 1.2 µg/mL before saturation of SAv binding sites with biotinylated GAGs at 10 µg/mL for 40 min. BMP2 was allowed to bind to GAGs with a 90-minute incubation step at 0.02 µg/mL (0.768 nM) in Hepes supplemented with 0.02% Tween^®^ 20 (Hepes-T) to prevent aggregation at physiological pH during the process. To remove the non-adsorbed BMP2, two rinsing steps were performed in Hepes-T and three rinsing steps in Hepes before cell seeding.

When preparing samples for automated immunofluorescence (IF) analysis, gold-coated surfaces were attached to the bottom side of a bottomless 96-well plate (Greiner Bio-one) using double-sided adhesive tape (FRAP Sandwich set, Paul Marienfeld), dividing one surface into four wells.

Glass biomimetic surfaces were built on glass-bottom 96-well plates (Greiner Bio-One) *via* a PLL(20)-g[3.5]-PEG(2)/PEGbiotin(3.4)50% linker (PLLgPEG, SuSoS AG). As previously described, SAv and cRGD were premixed (molar ratio 3:4) before incubation on the linker to allow an increased number of available biotin binding sites for the subsequent functionalization of GAGs^16^.

For both gold and glass platforms, the robot finally incubated BMP2 with adequate buffer and concentrations, indicated for each experiment. BMP2 was incubated between 30 and 90 min for cell experiments and 30 min for BMP2 immunofluorescence assays. When using 0.02% Tween-20 for BMP2 incubation, the unbound BMP2 was rinsed twice in Hepes-T and three rinsing steps in Hepes before cell seeding.

### Blocking steps on biomimetic platforms

A 2-hour blocking step was possibly performed to prevent BMP2 non-specific adsorption. The blocking was performed after SAv functionalization for gold platforms and PLLgPEG or SAv-RGD functionalization for glass platforms. The blocking step was performed using one of the following conditions: bovine serum albumin (BSA) at 3%, 5% or 10% (BP9703, ThermoFisher Scientific), D-(+)-Trehalose at 0.6 M (A19434.14, ThermoFisher Scientific), BSA/Trehalose mix (10%/0.6M), myoglobin at 1 mg/mL (M1882, Sigma-Aldrich), m-PEG2-NHS ester (284BP-23656, Tebubio), imidazole at 50mM (Sigma-Aldrich), glycine at 0.2 M (Sigma-Aldrich), and arginine at 0.2 M (Sigma-Aldrich). After blocking, the biomimetic platforms were rinsed 5 times with Hepes.

### *Ex-situ* characterization of BMP2 binding with quartz crystal microbalance

We measured with quartz crystal microbalance with dissipation monitoring (QCM-D, QSense Analyzer, Biolin Scientific) the shifts in resonance frequency (Δf in Hz) and energy dissipation (ΔD in dissipation units, ppm=10^−6^) to characterize the binding events in the biomimetic platforms sequential buildup. Experiments were performed at 24°C, and Δf and ΔD were measured at six overtones (i = 3, 5, …, 13)^12^. Only dissipation and normalized frequency of the third overtone (i = 3) are presented. Frequency measurements are related to the hydrated mass bound on the surface, while dissipation measurements are associated with the rigidity of the deposed molecule film. Platforms were functionalized on silicon dioxide (SiO2) crystals (QSX303, Biolin Scientific) as previously described^16^. BMP2 was injected at a 5 µg/mL concentration in Hepes buffer on crystals functionalized with SAv and cRGD, with or without HS. For BMP2, we used a “fast injection” procedure to mimic the functionalization performed by our liquid-handling robot. The solution of BMP2 was injected at a high flow rate of 100 µL/min until the liquid chamber was filled. The flow was stopped at this moment to let the molecules bind in a static regime. All measurements of frequency and dissipation shifts were performed after stabilizing the signal.

### Cellular assays

C2C12 cells were acquired from the ATCC (CRL1772) and cultured in DMEM with high glucose, pyruvate and GlutaMAX^™^ Supplement (cat. 10569010, Gibco^™^). Wild-type CHO-K1 cell lines (CHO-K1: CCL-61^™^) were cultured in DMEM/F-12 GlutaMAX (cat. 10565018, Gibco™). Both cell lines were cultured below the confluence, at 37°C under a 5% CO2 atmosphere, in polystyrene cell culture flasks (Falcon^®^, Corning) using the previously indicated media supplemented with 10% heat-inactivated fetal bovine serum (FBS, cat. 10270-106, Gibco™), and 1% of antibiotic-antimycotic (cat. 15240062, Gibco™). Cells were discarded after reaching a passage number of 12. Cells were serum-starved for 4 hours, detached with Accutase (A6964, Sigma-Aldrich), and plated on functionalized surfaces in 96-well plates. Each condition involved seeding 10,000 cells in coated wells of the 96-well plates maintained at 37°C with 5% CO2. Positive and negative control corresponds to cRGD platforms with or without 0.1 µg/mL of soluble BMP2 (sBMP2). The effect of BMP2 adsorbed on GAGs or non-specifically was studied by rinsing unbound BMP2 before cell seeding. Cells were fixed after 90 min using 2% PFA for 5 min, then 4% PFA for 20 min. Each condition was assessed with intra-experimental technical duplicates.

### Immunofluorescence assays on biomimetic platforms

A standard ELISA assay was performed to detect the binding of BMP2 on the platforms. After a 1-hour blocking step with 3% BSA in PBS, a polyclonal anti-BMP2/BMP4 primary antibody (AF355, R&D Systems) was incubated at 10 µg/mL for 2 h with 3% BSA in PBS at room temperature. Samples were washed with PBS, incubated with a secondary Cy3-conjugated donkey anti-mouse antibody (Jackson ImmunoResearch) for 1 h in 3% BSA, and washed. Mean fluorescence intensity at 595±35 nm was measured with a spectrophotometer microplate reader (Spark^®^, Tecan) with 535±25 nm wavelength excitation. Measurements of technical duplicates were plotted as mean ± standard deviation (SD).

For cellular assays, we adapted existing protocols already published to quantify the nuclear translocation of pSMAD1/5/9 via immunofluorescence (IF)^16, 28^. Briefly, fixed cells were first permeabilized with 0.2% Triton X-100 (Sigma-Aldrich) (w/v) before following the same steps as indicated above (blocking, primary antibody, fluorescent-conjugated secondary antibody). Primary rabbit anti-pSMAD1/5/9 (Cell Signaling Technology) was diluted at 1:400, and the secondary antibody (1:500, goat anti-rabbit Alexa Fluor 488, Thermo Fischer Scientific) was incubated simultaneously with rhodamine-phalloidin (1:1000, Thermo Fischer Scientific) and DAPI (1:1000, Sigma-Aldrich). Cell imaging was performed utilizing the InCell Analyzer 2500 (GE Healthcare) microscope with the 20x objective across three channels. Following established protocols, subsequent image analysis was conducted using the automated software InCarta (Molecular Devices). The intensity of pSMAD1/5/9 was evaluated exclusively within the nucleus, employing a mask derived from the DAPI signal, and the measurements were background-corrected for a minimum of 100 cells per well.

### Alkaline phosphatase staining

For ALP staining, we used fast blue RR salt (Sigma-Aldrich) within a 0.01% (w/v) naphthol AS-MX solution (Sigma Aldrich), following the manufacturer’s guidelines. ALP staining was quantified under liquid conditions by capturing images of the 96-well plate with the 4x objective of the InCell Analyzer 2500 microscope in brightfield mode. The acquired brightfield images were processed using ImageJ, wherein a low-intensity threshold was applied to assess the positive ALP area per well.

## Results

### Low BMP2 non-specific binding on the biomimetic SAv platforms induces strong cell signaling

Previous studies demonstrated with QCM-D that gold-based biomimetic SAv platforms were well passivated against the non-specific binding of BMP2^11^. Therefore, we employed them to investigate the influence of GAGs on BMP signaling by evaluating nuclear translocation of pSMAD1/5/9 in C2C12 cells seeded on gold biomimetic platforms. We adsorbed BMP2 at a 0.1 µg/mL concentration to different GAGs and rinsed the unbound proteins before seeding cells. Following a 90-minute incubation period on the platforms, we quantified pSMAD1/5/9 within cell nuclei *via* high-content immunofluorescence analysis. Strikingly, as illustrated in **Figure 1A**, the adsorbed BMP2 led to similar increases in pSMAD1/5/9 regardless of the immobilized GAG on the platform. This outcome was unexpected, given the significant differences in affinity we have previously observed for BMP2 interactions with distinct GAGs.

**Fig. 1;.**
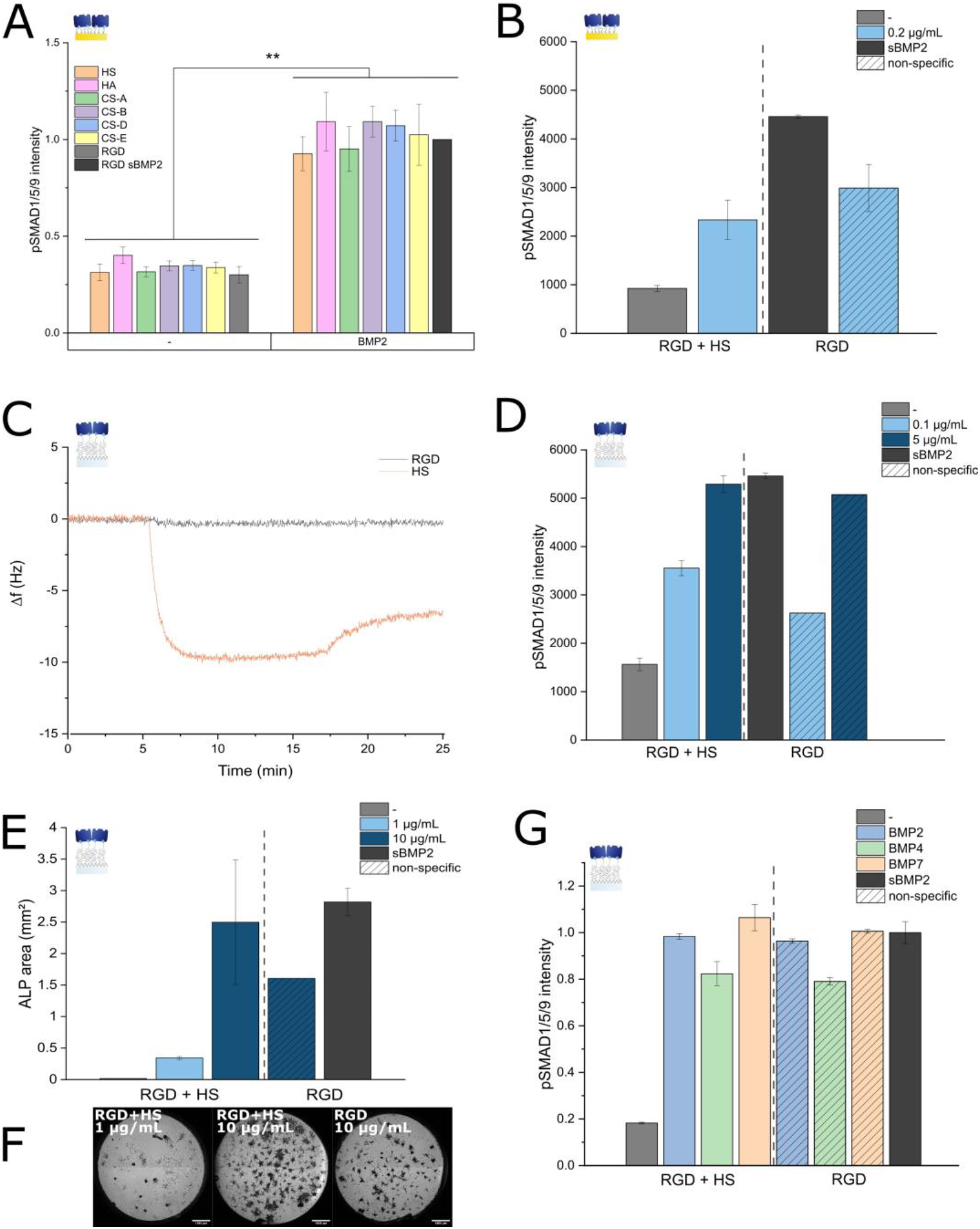
BMP2 binds non specifically to biomimetic platforms. For graphs **A, B, D, E, G**, positive (black) and negative (gray) controls correspond to cRGD functionalized platforms (without HS) exposed to soluble BMP2 at 0.1µg/mL or not. A. Mean fluorescence intensity of nuclear pSMAD1/5/9 in C2C12 cells after a 90-minute culture on gold biomimetic platforms with or without BMP2. BMP2 was either incubated at 0.2 µg/mL with GAGs followed, or added directly in solution with cells at 0.1µg/mL (sBMP2). Mean values were normalized by the positive sBMP2 positive control ± SEM (n=3). **B. D**. Mean fluorescence intensity of nuclear pSMAD1/5/9 in C2C12 cells after a 90-minute culture on **B**. gold or **D**. glass biomimetic platforms with or without BMP2. The HS platforms were incubated with 0.1, 0.2 or 5.0 µg/mL BMP2. cRGD platforms without GAGs correspond to the non-specific cellular response. Mean fluorescence values are plotted ± SD C. Frequency shifts measured via QCM-D upon binding of BMP2 at 5µg/mL on glass biomimetic platforms, specifically on HS, or non-specifically on platforms with PLLgPEG, SAv and RGD only. **E**. Alkaline phosphatase (ALP) positive area after 3 days of culture on glass biomimetic platforms. The platforms harboring HS were incubated with 1 (light blue) or 10 µg/mL (dark blue) BMP2. Cellular response induced by BMP2 non-specific binding is represented with diagonal stripes. Mean ALP positive surface is plotted ± SD. **F**. Brightfield images of the entire wells with ALP accumulation in black. Scale bar is 1000 µm. **G**. Mean fluorescence intensity of nuclear pSMAD1/5/9 in C2C12 cells after a 90-minute culture on glass biomimetic platforms, with or without BMPs: BMP2 (blue), BMP4 (green) and BMP7 (beige). All BMPs were incubated at 1.0 µg/mL. Data are plotted as normalized mean fluorescence intensity ± SD.

In light of this, we conducted a control cellular experiment, comparing the effects of BMP2 incubation on RGD-functionalized gold biomimetic platforms with and without heparan sulfate (HS), a GAG exhibiting a high affinity for BMP2 (**Figure 1B**). This experiment revealed a slightly higher pSMAD1/5/9 level on platforms lacking HS, indicating a significant bioactivity even in the absence of non-specifically bound BMP2, as determined by QCM. This observation may explain why the signaling observed in **Figure 1A** was comparable across different GAGs, as the non-specific signaling falls within the same range as the specific signaling induced by HS-bound BMP2.

We next investigated if the non-specific binding of BMP2 remained an issue with the alternative construction of biomimetic platforms on glass substrates. We first characterized *via* QCM-D the binding of BMP2 on glass biomimetic platforms. Our results indicated that there is no non-specific binding of BMP2 on the PLL-g-PEG coated platforms as proved in Sefkow-Werner et al 2022^16^, but that BMP2 binds to immobilized HS (**Figure 1C**). We then plated C2C12 cells on biomimetic glass platforms to assess the impact of non-specific binding to RGD-functionalized platforms on pSMAD1/5/9 cellular response. In **Figure 1D**, we show that similarly to gold substrate platforms, non-specific signaling is in the same range as signaling mediated by HS-bound BMP2. Furthermore, we tested two different BMP2 concentrations and showed that the non-specific signaling is also dose-dependent, likely due to increased BMP2 adsorption at higher concentrations. The pSMAD1/5/9 level was an early marker of osteogenic differentiation, so we then explored whether the non-specific binding of BMP2 also influenced cellular signaling over an extended culture period. C2C12 cells were cultured on glass biomimetic platforms for 3 days before comparing their expression of alkaline phosphatase (ALP) as a marker of bone differentiation. We show in **Figure 1E-F** that for 10 µg/mL BMP2 concentration, the ALP expression was slightly lower on the RGD platform yet comparable to the ALP expression on HS platforms.

To explore if this non-specific binding was a unique feature of BMP2, we conducted a cellular experiment with IF analysis of nuclear pSMAD1/5/9 with other protein members of the BMP family, namely BMP4 and BMP7. As shown in **Figure 1G**, for all BMPs, the non-specific signal induced by BMPs adsorbed on RGD-functionalized glass platforms was always comparable to the signal on HS-functionalized platforms. BMP2 purchased from another provider (R&D Systems) also induced non-specific signaling (data not shown).

Overall, the non-specific binding of BMP2 on glass or gold biomimetic platforms, even if not detectable with QCM-D, triggers a signaling response. The non-specific signal induced by adsorbed BMP2 was also dose-dependent and sustained for at least three days, as observed with ALP staining. The non-specific binding of BMP2 is a characteristic shared with other members of the BMP family, such as BMP4 and BMP7.

### BMP2 aggregates at physiological pH, but the native state can be rescued by Tween-20

The literature shows that BMP2 tends to aggregate under physiological pH conditions^29, 30^. Considering that aggregation may contribute to significant non-specific binding, we explored whether using adequate buffer conditions to avoid BMP2 aggregation could prevent its non-specific adsorption. Given that the isoelectric point (pI) of BMP2 is 8.2 ± 0.4, its solubility is greatly improved at acidic pH^30^. To allow its interaction with negatively charged sulfate groups of GAGs, the pH must be below the pI so that BMP2 is positively charged. In our research of BMP2 aggregation-free conditions, we used dynamic light scattering (DLS) to measure the size distribution of BMP2 particles in different buffers (**Figure. 2A-F**). In the recommended HCl stock buffer at pH 3.0, the mean particle size (hydrodynamic diameter) measured 5.56 ± 1.39 nm, close to previously reported values for BMP2 in MES buffer at pH 5.0 with a diameter of 7.8 ± 0.7 nm^29^. In contrast, as expected, BMP2 formed aggregates in Hepes and PBS buffers at pH 7.4, larger than 100 nm or 500 nm for Hepes and PBS, respectively. Interestingly, adding Tween-20 surfactant into the buffers rescued the native state of BMP2 observed in the HCl buffer, with a mean particle size of 7.62 ± 0.4 nm and 6.87 ± 0.67 nm in Hepes-T and PBS-T, respectively. Alternatively, using below-neutral pH at 4.2 with an acetate buffer also conserved the initial state of BMP2 with a particle size of 5.24 ± 0.12 nm. Hence, pH adjustments and using surfactants such as Tween-20 preserve the native state of BMP2 and prevent its aggregation. It is worth noting that the use of Hepes buffer without salt also improved the BMP2 solubility, with a particle size of 13.3 ±1.7 nm, which could be explained by complex salting-in and salting-out effects dependent on the salt concentration, as previously observed^31^. With the new conditions found to avoid BMP aggregation, we performed a cellular experiment on glass biomimetic platforms to investigate their effect on the non-specific BMP2 signaling. Unfortunately, the pSMAD1/5/9 level of C2C12 cells was still the same between HS and the non-specific conditions, both with Hepes-T or Acetate-T buffers, indicating that the BMP2 adsorption on platforms was not exclusively due to protein aggregation (**Figure. 2G**). Nevertheless, we consistently included 0.02% Tween-20 in buffers in subsequent experiments to prevent BMP2 aggregation.

**Figure. 2.**
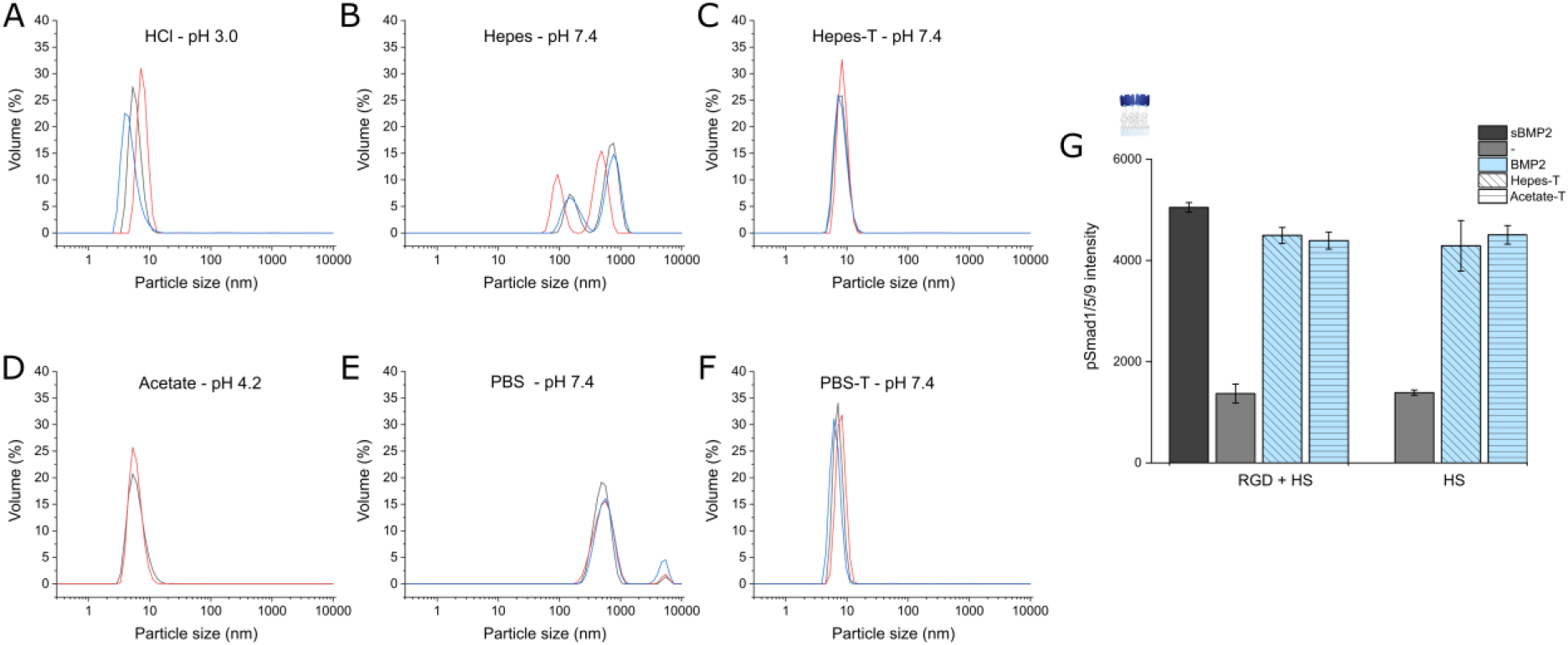
DLS size distribution by volume of BMP2 at 100µg/mL in different buffers: **A** HCl 1mM, pH 3.0; **B** Hepes, pH 7.4; **C** Hepes-T, pH 7.4; **D** Sodium acetate, pH 4.2; **E** PBS, pH 7.4; **F** PBS-T, pH 7.4. Triplicate measurements are represented in blue, black and orange. **G** Mean fluorescence intensity of nuclear pSMAD1/5/9 in C2C12 cells after a 90-minute culture on glass biomimetic platforms, with or without BMP2. BMP2 was incubated at 1 µg/mL in Hepes-T (diagonal stripes) or acetate-T buffer (horizontal stripes). Mean fluorescence values are plotted ± SD.

### BMP2 mainly binds non-specifically to the base substrate of biomimetic platforms

Given that BMP2 aggregation was not the primary reason for its non-specific adsorption, we next explored whether it could involve non-specific interactions with one of the components forming the biomimetic platforms. To this end, we assessed the BMP2 adsorption at different stages of the glass biomimetic platform construction using either a fluorescently labelled BMP2 (BMP2-rhodamine) or a specific polyclonal anti-BMP2 antibody (for the latter, microscopy images are shown in **Figure. 3A**).

**Figure. 3.**
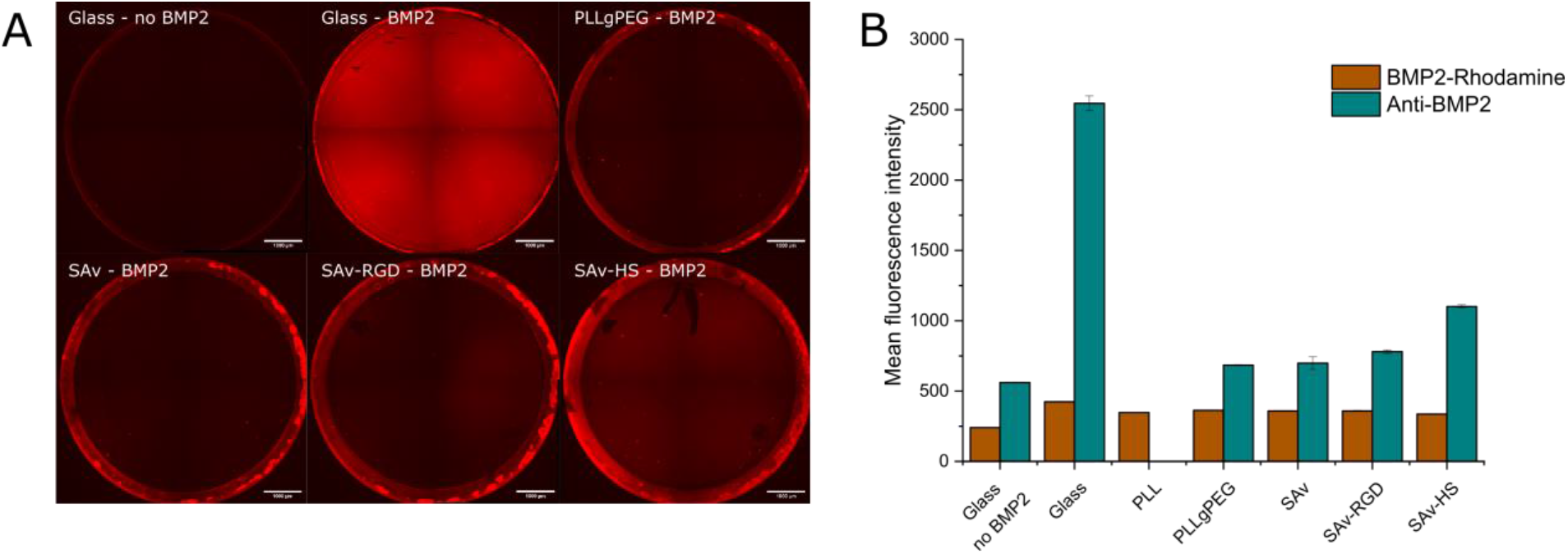
**A** Mean fluorescence intensity BMP2-Rhodamine or BMP2 detected with a specific antibody on glass biomimetic platforms after each functionalization step. Mean fluorescence values are plotted ± SD. **B** Representative fluorescence images of BMP2 binding detected via the anti-BMP2 antibody.

The protein adsorption was quantified with a fluorescence plate reader, which highlighted that the binding of both BMP2 and BMP2-rhodamine was most pronounced on the bare glass substrate (**Figure. 3B**). The functionalization with PLL or PLLgPEG reduced significantly the binding, although not to the extent of the negative control. The functionalization with SAv yielded no change in binding, and neither did RGD, except for a slight change detected with the BMP2 antibody. However, further analysis with BLI revealed no specific binding of BMP2 onto RGD (data not shown), suggesting that the observed variation was negligible. BMP2 bound effectively to HS, showing that even after rinsing steps, the amount of BMP2 seems to remain more important on the HS platforms than when non-specifically bound to the platforms.

In contrast to BMP2, the modified BMP2-rhodamine did not bind to HS and even showed a slightly reduced fluorescence. We indeed showed *via* QCM-D that the tagged BMP2 did not bind to HS (data not shown), undoubtedly due to the labelling procedure primarily affecting the amine groups of lysine residues, some of which are present in the N-terminal BMP2 domain responsible for HS interaction^32, 33^. Although the experiment was not conducted on gold platforms, the observed non-specific binding of BMP2 appears disconnected from the constituents of the platform (SAv, RGD). This implies that the predominant influence of non-specific BMP2 binding arises from the underlying substrate (glass or gold), which is insufficiently shielded by the PEG linkers. Increasing the grafting density of PEG chains should further reduce the binding. However, higher grafting densities of PEG on PLLgPEG copolymer products are not commercially available.

### Non-specific binding is electrostatic and can be somewhat reduced by several blocking strategies

Since the BMP2 adsorption arises from the underlying substrate of the biomimetic platforms (glass or gold), we attempted to reduce it by employing different blocking strategies to passivate the surfaces. We first tested the effect of various blocking molecules after the functionalization of SAv and RGD in a cell experiment, ensuring no interference with the initial steps of platform construction. As shown in **Figure 4A**, we blocked the glass biomimetic platforms with either 3% BSA, 1 mg/mL myoglobin or m-PEG2-NHS-Ester. While BSA is a common choice, we also assessed myoglobin, which is approximately fourfold smaller (16 kDa, approximately the size of BMP2) and could potentially infiltrate PEG films more easily to bind to non-specific sites on the glass surface. To ensure that BMP2 did not bind onto the PLL, we employed an m-PEG2-NHS-Ester linker to react with amine groups of PLL lysine residues. This approach could also increase the PEG density, potentially enhancing the platform’s anti-fouling properties.

**Figure 4.**
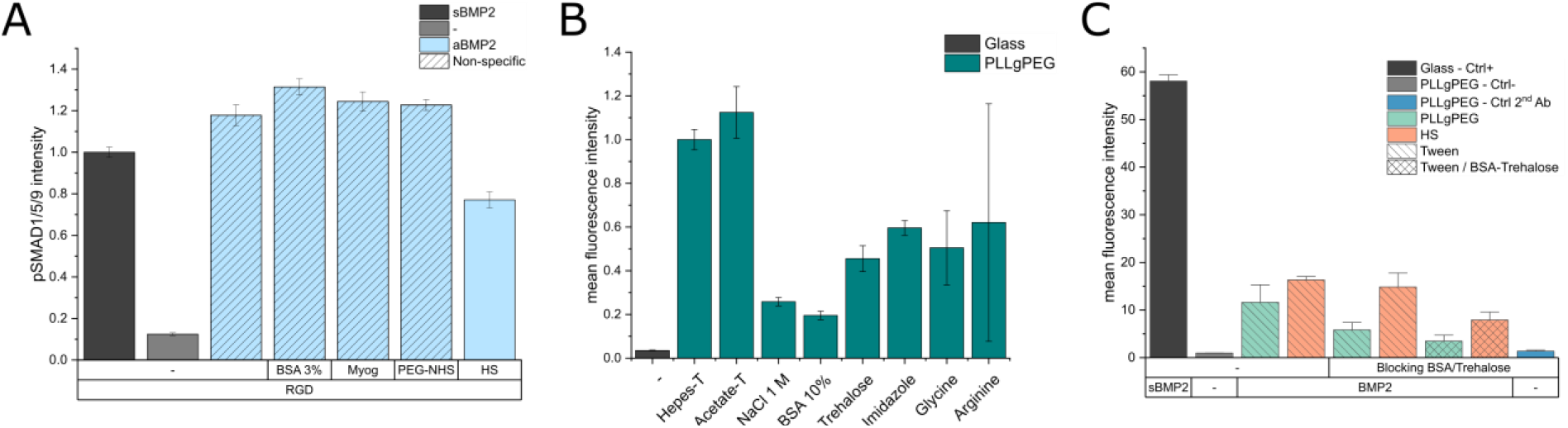
**A** Mean fluorescence intensity of nuclear pSMAD1/5/9 in C2C12 cells after a 90-minute culture on glass biomimetic platforms, including a blocking step with either 3% BSA, 1 mg/mL myoglobin or m-PEG2-NHS-Ester. BMP2 was incubated at 1 µg/mL in Hepes-T. Mean fluorescence values are plotted ± SD. **B** Mean fluorescence intensity of BMP2 detected with a specific antibody on PLLgPEG-functionalized glass, using different reagents for the blocking step and supplemented with Tween-20 for the BMP2 incubation buffer. The negative control without BMP2 on the glass is grey. Mean fluorescence values were normalized to the Hepes-T condition on PLLgPEG and plotted ± SD. **C** Mean fluorescence intensity of BMP2 detected with a specific antibody. Intensity comparison on glass biomimetic platforms with or without HS, with different combinations of blocking and BMP2 incubation buffers: 1) No blocking and BMP2 in Hepes-T; 2) BSA/Trehalose blocking and BMP2 in Hepes-T; 3) BSA/Trehalose blocking and BMP2 in Hepes-T with BSA/Trehalose. BMP2 was incubated on platforms at five µg/mL. A secondary antibody control on BSA-Trehalose blocked platforms were included (blue). Mean fluorescence values are plotted ± SEM for at least two experiments.

However, none of these options sufficiently reduced the non-specific binding since the pSMAD1/5/9 signaling was even higher than on HS signaling platforms. Comparable outcomes were observed when using Poloxamer 188 (also known as Pluronic^®^ F-108), a non-ionic surfactant frequently employed in biological formulations. Poloxamer 188 is a tri-block copolymer of polyethylene oxide – polypropylene oxide – polyethylene oxide (PEO–PPO– PEO). It usually binds surfaces through its central PPO hydrophobic part while the two PEO hydrophilic tails extend in solution. Self-assembled monolayers of Poloxamer 188 have demonstrated good anti-fouling properties^34^. However, probably due to restrained surfactant adsorption influenced by PEG chains on the platforms (QCM-D data not shown), the surfactant showed no effect on the non-specific signaling in our cellular assay.

For the following tests, we investigated with the anti-BMP2 antibody the efficacy of various conditions for passivating the surfaces; indeed, SMAD 1/5/9 is rapidly phosphorylated in the presence of the minimal amount of BMP2, preventing sensitivity to small improvements of the passivation. As shown in **Figure 4B**, we next tested an expended range of reagents at high concentrations to prevent the binding of BMP2 on the platforms. Each reagent was used in the blocking step and the BMP2 incubation buffer. The binding was quantified *via* IF by the anti-BMP2 antibody. Remarkably, we observed that the non-specific binding was significantly reduced using 1 M of NaCl, indicating that it relies, at least in part, on electrostatic interactions. However, the use of high salt in our experimental setup could not be considered, as the binding of BMP2 to GAGs also involves electrostatic interactions. We also showed that increasing the BSA concentration to 10% reduced the non-specific binding by about 80%, similarly to the 1 M of NaCl condition. Trehalose, imidazole, glycine, and arginine were other compounds tested for blocking non-specific binding. All exhibited a moderately positive effect, reducing BMP2 adsorption on PLLgPEG by approximately 50%. Among these, trehalose displayed a slightly better effect than the other molecules and was thus used in combination with 10% BSA for the blocking step.

As shown in **Figure 4C**, performing a blocking step with 10% BSA/0.6 M trehalose reduced the non-specific binding of BMP2 on PLLgPEG without affecting its binding to HS. The comparably lesser effect of the BSA and trehalose combination compared to the experiment shown in **Figure 4B** could be attributed to inherent variability in non-specific binding across different experiments. In contrast, combining the blocking step with incubation of BMP2 in a Hepes-T buffer containing 10% BSA and 0.6 M trehalose further reduced the non-specific binding and concurrently lowered the specific binding to HS. Overall, the precise contributions of BSA and trehalose in the blocking step or within the BMP2 incubation buffer remain unclear. Still, their combination as blocking agents undoubtedly reduced the non-specific binding.

### Gold-based biomimetic platforms prevent better BMP2 non-specific signals with respect to glass-based platforms

To explore the effects of BSA and trehalose in reducing the BMP2 non-specific signaling, we performed cellular experiments on gold and glass platforms with C2C12 and CHO cells, using two different blocking formulations: either 5% BSA alone or 10% BSA with 0.6 M trehalose. We show in **Figure 5A-B** the comparison of the mean nuclear pSMAD1/5/9 intensity between the non-specific signal (RGD) and the signal mediated by HS platforms (RGD+HS), considering distinct platform substrates and passivation conditions. If the basal non-specific signaling seemed to be reduced for CHO cells compared to C2C12 cells, there was a consistent reduction of non-specific signaling on gold platforms compared to glass platforms, for both cell types. Indeed, while the surface passivation with 5% BSA on the glass platform only slightly reduced the non-specific signal, the reduction amounted to approximately 50% on gold platforms. In comparison to BSA at 5%, the application of the BSA/trehalose mix reduced slightly more the C2C12 non-specific signal on glass platforms (**Figure 5A**) and reduced it significantly on gold platforms for CHO cells (**Figure 5B**). Surprisingly, BSA/trehalose blocking on gold platforms with C2C12 cells led to a strikingly elevated pSMAD1/5/9 signaling. Overall, the gold platforms provide better anti-fouling properties than glass platforms, probably due to a more organized layer or PEG chains. Additionally, CHO cells apparently displayed reduced responsiveness to non-specifically bound BMP2 compared to C2C12 cells, making them potentially more appropriate cellular models for limiting the impact of non-specific signaling.

**Figure 5.**
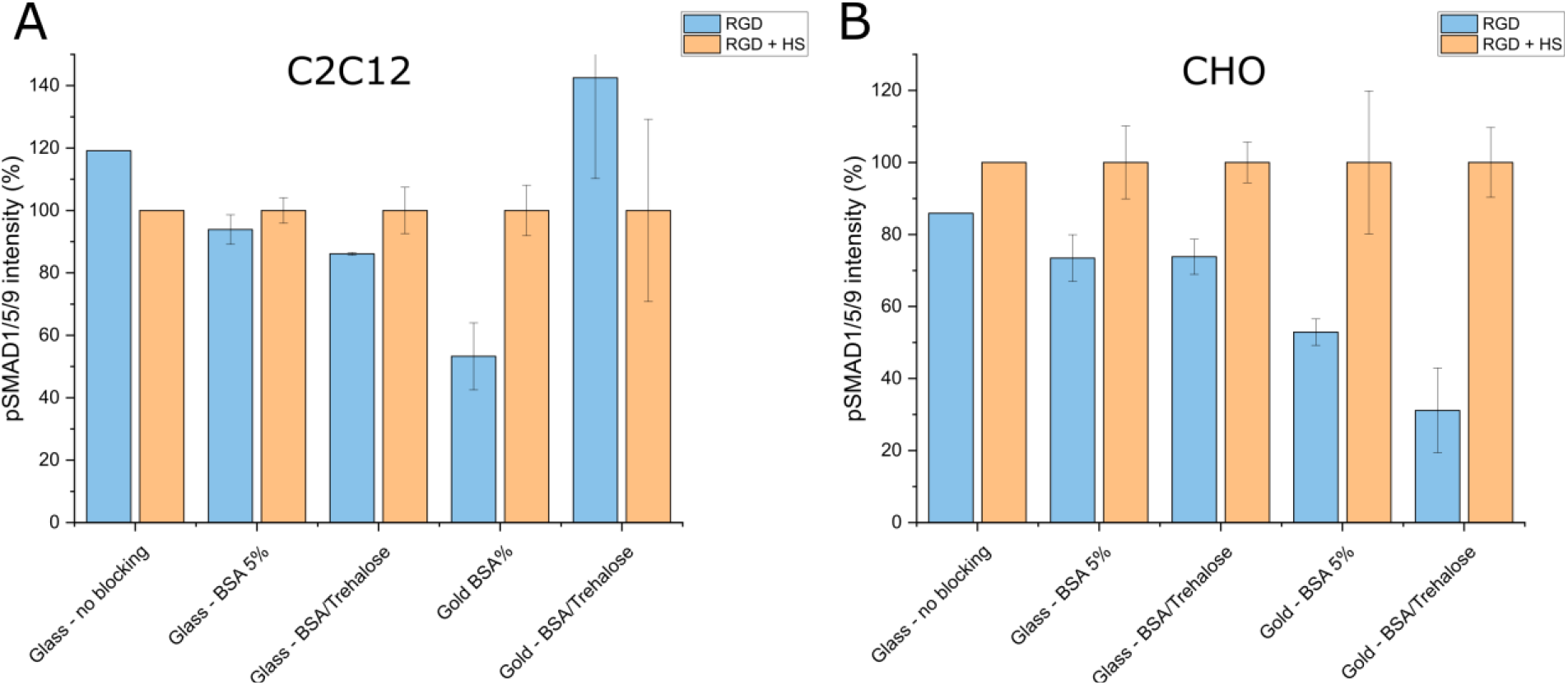
Mean fluorescence intensity of nuclear pSMAD1/5/9 in **A** C2C12 or **B** CHO cells after a 90-minute culture on glass or gold biomimetic platforms functionalized, including a blocking step with 5% BSA or BSA/Trehalose (10%/0.6M) before BMP2 incubation. Mean fluorescence values are normalized to each HS condition ± SD.

## Discussion

Our study highlights that moderate protein non-specific binding on biomaterials (here, BMP2 on biomimetic platforms) can trigger an important cell signaling response. The non-specific binding of proteins on biomaterials should be considered in their design and for applications – both *in vitro* and *in vivo* – to correctly interpret the effect of biomaterials on the bioactivity of proteins. During our investigations, the BMP2-induced non-specific signaling initially prevented us from elucidating the distinct roles of GAGs in BMP2 signaling regulation. We established that this non-specific adsorption (and signaling) was dose-dependent, relatively stable, bioactive for up to three days, and was present independently of the biomimetic platform substrate (glass or gold). This issue was also extended to other BMP family proteins, including BMP4 and BMP7. We determined that the primary cause of non-specific binding is the platform’s underlying substrate (glass or gold) rather than its constituents like SAv or RGD. We next explored conditions susceptible to reducing the non-specific signaling to tackle this issue. Addressing this challenge was complex, with the objectives of retaining the GAG-binding capacity of BMP2, preserving the substrate transparency for high-throughput microscopy, and avoiding interference with the platform construction.

We investigated the BMP2 aggregation in different media and confirmed that BMP2 aggregates at physiological pH but can be solubilized with acidic pH buffers^29, 30^. Remarkably, we discovered that adding surfactants such as Tween-20 could prevent BMP2 aggregation, even at neutral pH. Of note, the use of Hepes buffer without salt enhanced BMP2 stability, likely due to complex “salting-out” and “salting-in” effects. As previously observed, the addition of NaCl in MES buffer (pH 5) was demonstrated to cause BMP2 precipitation at concentrations of only 80 mM NaCl (salting-out)^31^. Further increasing the NaCl concentration above 0.3 M increased the solubility (salting-in) until decreasing again above 1 M.

Our research revealed that BMP2 non-specific adsorption relies on electrostatic interactions that could be disrupted with high salt concentrations, though incompatible with GAG-BMP2 interactions. We then showed that some blocking reagents at high concentrations, particularly BSA and trehalose, could reduce the BMP2 non-specific binding by 50 % to 80 %. Although this study did not focus on clarifying the specific contributions of BSA and trehalose as blocking agents or additives in the BMP2 buffer formulation, their combined use as blocking agents unequivocally led to an important reduction of the non-specific binding.

While BSA is commonly used in immunofluorescence both for the blocking step and as an additive in the incubation of antibodies, trehalose has been primarily used for biological formulations^35, 36^. Trehalose, recognized for preventing protein aggregation, has been studied in particular for treating neurodegenerative diseases, such as Alzheimer’s and Huntington’s^37^. It also showed some interesting properties in BioLayer Interferometry studies as an additive for reducing non-specific binding^38^. Interestingly, a patent described using 5% trehalose to block functionalized surfaces, reducing serum protein adsorption^39^.

Our findings suggested that CHO cells might be less sensitive to non-specifically bound BMP2 than C2C12 cells. We also determined that gold platforms exhibited superior anti-fouling properties under identical blocking conditions compared to glass. The distinct PEG-based linkers for the different substrates may contribute to these differences. Therefore, out of the scope of this study, another option to reduce the non-specific signaling could be to design alternative constructions of the platforms with improved anti-fouling properties. Potential strategies include functionalizing poloxamers conjugated with an appropriate amount of biotin to build the streptavidin monolayer. Other strategies inspired by cell membrane properties could be considered, such as the functionalization of phosphorylcholines (PC) exhibiting non-fouling properties^40, 41^. Studies on protein adsorption with PC phospholipid films underscored their strong resistance to protein adsorption^12, 42, 43^. Preferably, platform development should be achieved on optically transparent surfaces for compatibility with established high-throughput functionalization and readout strategies. In summary, the findings of this study have implications for the design of biomaterials intended for investigating cellular responses upon bound proteins. Despite a relatively low non-specific protein binding detected *via* biophysical surface characterization techniques, it does not exclude a potentially significant impact at the cellular level. This study also provides new information about BMP2 aggregation and adsorption at surfaces. We show, in particular, that the combination of BSA and trehalose exhibits good performance as a blocking agent. Although the significant bioactivity of non-specifically bound BMP2 was an issue in our case, this study demonstrated the potential of biomaterials with bare glass in effectively retaining and delivering biologically active BMP2, as previously investigated^[38]^.

## Acknowledgements

We want to thank Marianne Weidenhaupt and Franz Bruckert for their fruitful discussions and the use of the DLS equipment at the LMGP Grenoble. We thank also Cesar Rodriguez-Emmenegger and Jenny Englert for the interesting discussions on this issue of non-specific binding of BMP2. We acknowledge the PTA facility, Upstream Technological Platform, for the use of the evaporator equipment. This work used the platforms of the Grenoble Instruct-ERIC center (ISBG; UAR 3518 CNRS-CEA-UGA-EMBL) within the Grenoble Partnership for Structural Biology (PSB), supported by FRISBI (ANR-10-INBS-0005-02) and GRAL, financed within the University Grenoble Alpes graduate school (Ecoles Universitaires de Recherche) CBH-EUR-GS (ANR-17-EURE-0003). Authors acknowledge the BLI platform scientific responsible, Jean-Baptiste REISER PhD, for its help and assistance. This work was funded by the ANR (GlyCON, grant number [ANR-19-CE13-0031-01 PRCI] and by the “Investissements d’avenir” program Glyco@Alps, grant number [ANR-15-IDEX-02]. This work has been supported by CNRS GDR 2088 “BIOMIM”, ANR-17-EURE-0003 and GRAL. IBS (Institut de Biologie Strucutrale) acknowledges integration into the IRIG (CEA).

## References

1. Silva-Bermudez, P.; Rodil, S. E., An overview of protein adsorption on metal oxide coatings for biomedical implants. Surface and Coatings Technology 2013, 233, 147–158.

2. Lichtenberg, J. Y.; Ling, Y.; Kim, S., Non-Specific Adsorption Reduction Methods in Biosensing. Sensors (Basel, Switzerland) 2019, 19 (11), 2488.

3. Andrade, J. D.; Hlady, V. In Protein adsorption and materials biocompatibility: A tutorial review and suggested hypotheses, 1986; Springer: 1986; pp 1–63.

4. Schlenoff, J.; B., Zwitteration: Coating Surfaces with Zwitterionic Functionality to Reduce Nonspecific Adsorption. Langmuir : the ACS journal of surfaces and colloids 2014, 30 (32), 9625–9636.

5. Otsuka, H.; Nagasaki, Y.; Kataoka, K., Self-assembly of poly(ethylene glycol)-based block copolymers for biomedical applications. Current Opinion in Colloid & Interface Science 2001, 6 (1), 3–10.

6. Ngo, B. K. D.; Grunlan, M. A., Protein Resistant Polymeric Biomaterials. ACS Macro Letters 2017, 6 (9), 992–1000.

7. Lowe, S.; O’Brien-Simpson, N. M.; Connal, L. A., Antibiofouling polymer interfaces: poly(ethylene glycol) and other promising candidates. Polymer Chemistry 2014, 6 (2), 198-212.

8. Xiao, X.-F.; Jiang, X.-Q.; Zhou, L.-J., Surface Modification of Poly Ethylene Glycol to Resist Nonspecific Adsorption of Proteins. Chinese Journal of Analytical Chemistry 2013, 41 (3), 445–453.

9. Michel, R.; Pasche, S.; Textor, M.; Castner, D. G., The Influence of PEG Architecture on Protein Adsorption and Conformation. Langmuir : the ACS journal of surfaces and colloids 2005, 21 (26), 12327–12332.

10. Li, L.; Chen, S.; Jiang, S., Protein interactions with oligo(ethylene glycol) (OEG) self-assembled monolayers: OEG stability, surface packing density and protein adsorption. Journal of Biomaterials Science, Polymer Edition 2007, 18 (11), 1415–1427.

11. Migliorini, E.; Horn, P.; Haraszti, T.; Wegner, S.; Hiepen, C.; Knaus, P.; Richter, P.; Cavalcanti-Adam, E., Enhanced biological activity of BMP-2 bound to surface-grafted heparan sulfate. Advanced Biosystems 2017, 1 (4), 1600041.

12. Migliorini, E.; Thakar, D.; Sadir, R.; Pleiner, T.; Baleux, F.; Lortat-Jacob, H.; Coche-Guerente, L.; Richter, R. P., Well-defined biomimetic surfaces to characterize glycosaminoglycan-mediated interactions on the molecular, supramolecular and cellular levels. Biomaterials 2014, 35 (32), 8903–15.

13. Sefkow-Werner, J.; Machillot, P.; Sales, A.; Castro-Ramirez, E.; Degardin, M.; Boturyn, D.; Cavalcanti-Adam, E. A.; Albiges-Rizo, C.; Picart, C.; Migliorini, E., Heparan sulfate co-immobilized with cRGD ligands and BMP2 on biomimetic platforms promotes BMP2-mediated osteogenic differentiation. Acta biomaterialia 2020.

14. Thakar, D.; Dalonneau, F.; Migliorini, E.; Lortat-Jacob, H.; Boturyn, D.; Albiges-Rizo, C.; Coche-Guerente, L.; Picart, C.; Richter, R. P., Binding of the chemokine CXCL12alpha to its natural extracellular matrix ligand heparan sulfate enables myoblast adhesion and facilitates cell motility. Biomaterials 2017, 123, 24–38.

15. Migliorini, E.; Horn, P.; Haraszti, T.; Wegner, S. V.; Hiepen, C.; Knaus, P.; Richter, R. P.; Cavalcanti-Adam, E. A., Enhanced Biological Activity of BMP-2 Bound to Surface-Grafted Heparan Sulfate. Advanced Biosystems 2017, 1 (4), 1600041–1600041.

16. Sefkow-Werner, J.; Le Pennec, J.; Machillot, P.; Ndayishimiye, B.; Castro-Ramirez, E.; Lopes, J.; Licitra, C.; Wang, I.; Delon, A.; Picart, C.; Migliorini, E., Automated Fabrication of Streptavidin-Based Self-assembled Materials for High-Content Analysis of Cellular Response to Growth Factors. ACS Applied Materials & Interfaces 2022, 14 (29), 34113–34125.

17. Sefkow-Werner, J.; Migliorini, E.; Picart, C.; Wahyuni, D.; Wang, I.; Delon, A., Combining Fluorescence Fluctuations and Photobleaching to Quantify Surface Density. Analytical chemistry 2022.

18. Marquetti, I.; Desai, S., Nanoscale Topographical Effects on the Adsorption Behavior of Bone Morphogenetic Protein-2 on Graphite. International Journal of Molecular Sciences 2022, 23 (5), 2432.

19. Marquetti, I.; Desai, S., Molecular modeling the adsorption behavior of bone morphogenetic protein-2 on hydrophobic and hydrophilic substrates. Chemical Physics Letters 2018, 706, 285–294.

20. Huang, B., Current molecular dynamics opinions on interactions between bone morphogenetic protein-2 and inorganic materials. Materials Today Communications 2023, 34, 105307.

21. Utesch, T.; Daminelli, G.; Mroginski, M. A., Molecular Dynamics Simulations of the Adsorption of Bone Morphogenetic Protein-2 on Surfaces with Medical Relevance. Langmuir : the ACS journal of surfaces and colloids 2011, 27 (21), 13144–13153.

22. Oliveira, A. F.; Gemming, S.; Seifert, G., Molecular dynamics simulations of BMP-2 adsorption on a hydrophobic surface. Materialwissenschaft und Werkstofftechnik 2010, 41 (12), 1048–1053.

23. Dong, X.; Wang, Q.; Wu, T.; Pan, H., Understanding Adsorption-Desorption Dynamics of BMP-2 on Hydroxyapatite (001) Surface. Biophysical journal 2007, 93 (3), 750–759.

24. Gilde, F.; Maniti, O.; Guillot, R.; Mano, J. F.; Logeart-Avramoglou, D.; Sailhan, F.; Picart, C., Secondary structure of rhBMP-2 in a protective biopolymeric carrier material. Biomacromolecules 2012, 13 (11), 3620–6.

25. Mulloy, B.; Gee, C.; Wheeler, S. F.; Wait, R.; Gray, E.; Barrowcliffe, T. W., Molecular weight measurements of low molecular weight heparins by gel permeation chromatography. Thrombosis and haemostasis 1997, 77 (4), 668–74.

26. Hamad, O. A.; Ekdahl, K. N.; Nilsson, P. H.; Andersson, J.; Magotti, P.; Lambris, J. D.; Nilsson, B., Complement activation triggered by chondroitin sulfate released by thrombin receptor-activated platelets. Journal of Thrombosis and Haemostasis 2008, 6 (8), 1413–1421.

27. Djerbal, L.; Vivès, R.; Lopin-Bon, C.; Richter, R.; Kwok, J. C. F.; Lortat-Jacob, H., Semaphorin 3A binding to chondroitin sulfate E enhances the biological activity of the protein, and cross-links and rigidifies glycosaminoglycan matrices. 2020.

28. Sales, A.; Khodr, V.; Machillot, P.; Chaar, L.; Fourel, L.; Guevara-Garcia, A.; Migliorini, E.; Albigès-Rizo, C.; Picart, C., Differential bioactivity of four BMP-family members as function of biomaterial stiffness. Biomaterials 2022, 121363.

29. Sundermann, J.; Zagst, H.; Kuntsche, J.; Wätzig, H.; Bunjes, H., Bone Morphogenetic Protein 2 (BMP-2) Aggregates Can be Solubilized by Albumin-Investigation of BMP-2 Aggregation by Light Scattering and Electrophoresis. Pharmaceutics 2020, 12 (12), E1143.

30. Crouzier, T.; Ren, K.; Nicolas, C.; Roy, C.; Picart, C., Layer-by-layer films as a biomimetic reservoir for rhBMP-2 delivery: controlled differentiation of myoblasts to osteoblasts. Small (Weinheim an der Bergstrasse, Germany) 2009, 5 (5), 598–608.

31. Quaas, B.; Burmeister, L.; Li, Z.; Satalov, A.; Behrens, P.; Hoffmann, A.; Rinas, U., Stability and Biological Activity of E. coli Derived Soluble and Precipitated Bone Morphogenetic Protein-2. Pharmaceutical research 2019, 36 (12), 184.

32. Ruppert, R.; Hoffmann, E.; Sebald, W., Human bone morphogenetic protein 2 contains a heparin-binding site which modifies its biological activity. European journal of biochemistry / FEBS 1996, 237 (1), 295–302.

33. Billings, P. C.; Yang, E.; Mundy, C.; Pacifici, M., Domains with highest heparan sulfate-binding affinity reside at opposite ends in BMP2/4 versus BMP5/6/7: Implications for function. The Journal of biological chemistry 2018, 293 (37), 14371–14383.

34. Chang, Y.; Chu, W.-L.; Chen, W.-Y.; Zheng, J.; Liu, L.; Ruaan, R.-C.; Higuchi, A., A systematic SPR study of human plasma protein adsorption behavior on the controlled surface packing of self-assembled poly(ethylene oxide) triblock copolymer surfaces. Journal of Biomedical Materials Research Part A 2010, 93A (1), 400–408.

35. Jain, N. K.; Roy, I., Effect of trehalose on protein structure. Protein science : a publication of the Protein Society 2009, 18 (1), 24–36.

36. Kaushik, J. K.; Bhat, R., Why Is Trehalose an Exceptional Protein Stabilizer?: AN ANALYSIS OF THE THERMAL STABILITY OF PROTEINS IN THE PRESENCE OF THE COMPATIBLE OSMOLYTE TREHALOSE *. Journal of Biological Chemistry 2003, 278 (29), 26458–26465.

37. Khalifeh, M.; Barreto, G. E.; Sahebkar, A., Therapeutic potential of trehalose in neurodegenerative diseases: the knowns and unknowns. Neural Regeneration Research 2021, 16 (10), 2026–2027.

38. Dubrow, A.; Zuniga, B.; Topo, E.; Cho, J.-H., Suppressing Nonspecific Binding in Biolayer Interferometry Experiments for Weak Ligand–Analyte Interactions. ACS Omega 2022, 7 (11), 9206–9211.

39. Jogikalmath, G. Method for blocking non-specific protein binding on a functionalized surface. WO2008055080A2, 2008-05-08, 2008.

40. Chen, S.; Zheng, J.; Li, L.; Jiang, S., Strong Resistance of Phosphorylcholine Self-Assembled Monolayers to Protein Adsorption: Insights into Nonfouling Properties of Zwitterionic Materials. Journal of the American Chemical Society 2005, 127 (41), 14473–14478.

41. Tegoulia, V. A.; Rao, W.; Kalambur, A. T.; Rabolt, J. F.; Cooper, S. L., Surface Properties, Fibrinogen Adsorption, and Cellular Interactions of a Novel Phosphorylcholine-Containing Self-Assembled Monolayer on Gold. Langmuir : the ACS journal of surfaces and colloids 2001, 17 (14), 4396–4404.

42. Richter, R.; Mukhopadhyay, A.; Brisson, A., Pathways of Lipid Vesicle Deposition on Solid Surfaces: A Combined QCM-D and AFM Study. Biophysical journal 2003, 85 (5), 3035–3047.

43. Vermette, P.; Gauvreau, V.; Pézolet, M.; Laroche, G., Albumin and fibrinogen adsorption onto phosphatidylcholine monolayers investigated by Fourier transform infrared spectroscopy. Colloids and Surfaces B: Biointerfaces 2003, 29 (4), 285–295.

44. Jurchenko, C.; Chang, Y.; Narui, Y.; Zhang, Y.; Salaita, K. S., Integrin-generated forces lead to streptavidin-biotin unbinding in cellular adhesions. Biophysical journal 2014, 106 (7), 1436–46.

